# Domain-general cognitive control processes in bilingual switching: evidence from midfrontal theta oscillations

**DOI:** 10.1101/2024.01.15.575665

**Authors:** Ningjing Cui, Vitoria Piai, Xiaochen Y. Zheng

## Abstract

Language control in bilingual speakers is thought to be implicated in effectively switching between languages, inhibiting the non-intended language, and continuously monitoring what to say and what has been said. It has been a matter of controversy concerning whether language control operates in a comparable manner to cognitive control processes in non-linguistic domains (domain-general) or if it is exclusive to language processing (domain-specific). As midfrontal theta oscillations have been considered as an index of cognitive control, examining whether a midfrontal theta effect is evident in tasks requiring bilingual control could bring new insights to the ongoing debate. To this end, we reanalysed the EEG data from two previous bilingual production studies where Dutch-English bilinguals named pictures based on colour cues. Specifically, we focused on three fundamental control processes in bilingual production: switching between languages, inhibition of the nontarget language, and monitoring of speech errors. Theta power increase was observed in switch trials compared to repeat trials, with a midfrontal scalp distribution. However, this midfrontal theta effect was absent in switch trials following a short sequence of same-language trials compared to a long sequence, suggesting a missing modulation of inhibitory control. Similarly, increased midfrontal theta power was observed when participants failed to switch to the intended language compared to correct responses. Altogether, these findings tentatively support the involvement of domain-general cognitive control mechanisms in bilingual switching.

## Introduction

In natural conversation, bilinguals appear to show remarkable flexibility in constantly speaking in one language and switching to another given the communicative situations without difficulty. In fact, this process is not as effortless as it seems, as both languages are simultaneously activated regardless of the language being spoken (Colomé, 2001; Costa et al., 1999; Hermans et al., 1998; Starreveld et al., 2014). This necessitates a process to restrict language processing to the target language and avoid the interference from the nontarget language, commonly referred to as language control.

One of the most commonly used experimental paradigms to investigate language control is cued *language switching* (Declerck & Philipp, 2015; Meuter & Allport, 1999; Sikora et al., 2016; Zheng et al., 2018a). The paradigm normally involves the task of naming presented items (e.g., pictures, digits) with visual cues indicating the language in which the items should be named. Bilinguals usually perform poorer on switch trials where they need to switch between languages, characterized by longer reaction times (RTs) and increased error rates, compared to repeat trials where they stay in the same language. This is known as the language-switch cost (Christoffels et al., 2007a; Costa & Santesteban, 2004; Jackson et al., 2001; Verhoef et al., 2009; Zheng et al., 2018b). The primary explanation for the language-switch cost pertains to the involvement of inhibitory control: the non-target language is inhibited, and this inhibition carries over to the next trial; switch trials require overcoming this inhibition and retrieving the suppressed language, making them more demanding than repeat trials (Declerck & Philipp, 2015; Green, 1998; Jackson et al., 2001; Liu et al., 2014; Philipp & Koch, 2009). Recent research has strived to further understand how *inhibitory control* evolves over time. By manipulating the number of repeat trials before transitioning to a switch (“run length”), Zheng et al. (2018b, 2020) observed faster responses during switches following a shorter sequence of repeat trials (i.e., a short run) compared to those following a longer sequence of repeat trials (i.e., a long run; see also Meuter & Allport, 1999; Kleinman & Gollan, 2018). This effect can be explained by decreased inhibition over time: with the repeated use of the same language, the inhibition of the non-target language decreases, making it easier to be overcome at a switch.

The control processes of bilingual production also involve the *monitoring of speech* (Acheson et al., 2012; Declerck et al., 2017b; Gollan et al., 2011; Hartsuiker, 2014). Despite being fluent in language switching, bilingual speakers still occasionally make language selection errors, wherein they say a word in the non-target language instead of the intended equivalent (e.g., saying the Dutch word “paraplu” instead of the English equivalent “umbrella”). These language selection errors are more frequently encountered on switch trials than repeat trials in the language switching task (Zheng et al., 2018a; 2018b).

The aforementioned three control processes during bilingual production, namely language switching, inhibitory control, and speech monitoring, parallel three key cognitive control processes in non-linguistic tasks: shifting between mental sets, inhibition of prepotent responses, and monitoring of actions (Alexander & Brown, 2010; Miyake et al., 2000). This raises the question of whether language control operates within broader cognitive control mechanisms (domain-general) or relies exclusively on language control mechanisms (domain-specific). Despite extensive research for decades, this topic still remains debated. Support for language control being domain-general primarily stems from similarities in behavioural and neural patterns observed during language and non-linguistic control. These similarities include matched switch-cost effect (Declerck et al., 2017a; Prior & Gollan, 2013; Timmermeister et al., 2020), activation of common brain regions (e.g., lateral prefrontal cortex, Abutalebi & Green, 2008; de Bruin et al., 2014; Hernandez et al., 2000; Wang et al., 2009; anterior cingulate cortex, Abutalebi et al., 2008, 2012; Christoffels et al., 2007b; Gauvin et al., 2016; Guo et al., 2011; Rossi et al., 2021; and presupplementary motor area; Christoffels et al., 2007b; de Bruin et al., 2014; Guo et al., 2011; Rossi et al., 2021), as well as the presence of similar event-related potential (ERP) signals (e.g., N2; Jackson et al., 2001; Jamadar et al., 2015; Kang et al., 2020; Verhoef et al., 2010; Zheng et al., 2020; and the event-related negativity; Coulter & Phillips, 2022; Zheng et al., 2018a).

However, there is evidence suggesting a unique language control mechanism, supporting the contrasting domain-specific perspective (e.g., Branzi et al., 2015, 2016; Calabria et al., 2012, 2015; Gray & Kiran, 2016; Magezi et al., 2012; Weissberger et al., 2012). For instance, this includes differences in the effects of aging (Weissberger et al., 2012), distinct ERP signatures (Magezi et al., 2012), and variations in the engagement of certain brain regions (i.e., the ACC and pre-SMA) for the two types of control (Branzi et al., 2015). More converging evidence is required to comprehend this issue by targeting different neural makers or using different approaches to characterise the language network (Fedorenko & Thompson-Schill, 2014).

A novel perspective to explore the connection between language and action control is by examining neural oscillations, which refer to the rhythmic electric activity that arises in the brain as a response to stimuli. The oscillatory activity of a substantial number of neurons, which can be observed through EEG recordings, serves as a fundamental basis for human cognition, perception, and behaviours by effectively coordinating communication over extensive brain networks (Helfrich & Knight, 2016; Thut et al., 2012). Neural oscillations encompass various frequency bands, with theta oscillations (4–8 Hz) over the midfrontal cortex being notably considered as a compelling candidate for the engagement of cognitive control (Cavanagh & Frank, 2014; Cohen & Donner, 2013; Rawls et al., 2020). Midfrontal theta is considered to originate from the bilateral medial frontal cortex and the ACC (e.g., Asada et al., 1999; Cohen et al., 2008). Increased midfrontal theta power has been observed when more top-down control is required across diverse cognitive domains (e.g., Cavanagh & Frank, 2014; Cavanagh et al., 2012; Cohen & Donner, 2013). In task switching, increased theta power was observed in switches as opposed to repeats, consistent with the RT switch cost effect (Cooper et al., 2019). Similarly, in response inhibition paradigms such as the Go/NoGo task, increased theta power was found when participants were required to withhold their response on the NoGo trials in comparison to the Go trials, in which they were asked to respond (Eisma et al., 2021; Nigbur et al., 2011). Increased theta power has also been observed in action monitoring, triggered after an error is committed, in contrast to correct responses (Cavanagh et al., 2012; Luu et al., 2004). Given the co-occurrence of midfrontal theta activity with the N2 and ERN components, both of which are linked to cognitive control, midfrontal theta has been often considered as the oscillatory counterpart to these components (Cavanagh & Frank, 2014).

Considering the strong link between midfrontal theta oscillations and cognitive control in the literature, examining theta activity in bilingual switching can add to the debate concerning the domain-general vs. domain-specific issue. Insofar, research on midfrontal theta oscillations in bilingual speech production has been scarce. Prior studies on theta oscillations in the language domain have primarily involved monolingual production and have consistently revealed increased midfrontal theta power in conditions demanding increased control (Krott et al., 2019; Piai et al., 2014; Shitova et al., 2017). More recently, midfrontal theta oscillations have been observed in bilinguals during picture naming in mixed-language contexts, which requires more control than single-language contexts (Liu et al., 2022). However, no study has specifically examined the role of midfrontal theta in the switching processes themselves.

To offer a novel perspective on the nature of language control as a domain-general or domain-specific process, this study explored the midfrontal theta oscillations in bilingual switching, with a specific focus on three language control processes: language switching, inhibitory control, and speech monitoring. Towards this goal, we reanalysed the EEG data obtained from two previous bilingual switching studies (Zheng et al., 2020; Zheng et al., 2018a). Both studies employed cued language switching paradigms with Dutch-English bilinguals naming pictures based on colour cues. We anticipated observing midfrontal theta oscillations in bilingual switching that are similar to those reported in non-linguistic control tasks. Specifically, theta power increase was anticipated in conditions requiring higher levels of control, namely: switch > repeat (language switching), switches after a short run > switches after a long run (inhibitory control), and language selection errors at a switch> correct responses (speech monitoring). Besides, we expected the theta effect to have a midfrontal distribution, presumably reflecting the involvement of the domain-general cognitive control network (e.g., ACC, PFC).

## Methods

The datasets analysed here were from two prior publications (Zheng et al., 2020; Zheng et al., 2018a). The two original studies adhered to the principles outlined in the Declaration of Helsinki and obtained approval from the Faculty Ethics Committee at Radboud University (ECSW2015-2311-349). The complete details of the materials and data collection procedures can be found in the original papers, while only a concise summary is provided here.

### Participants and Experimental Designs

Participants were two cohorts of Dutch native bilingual speakers (Zheng et al. 2020: *N* = 25, age 19–27 years, seven males, 18 females; Zheng et al. 2018a: *N* = 24, 19–30 years, five males, 19 females) who exhibited comparable English proficiency profiles. Participants were instructed to name pictures based on colour cues, where the colours red and yellow indicated naming in Dutch, while green and blue indicated naming in English. Both studies employed the same picture naming paradigms, albeit slight variations in their experimental designs (Figure 1). In order to probe the inhibitory control dynamics, one study experimentally varied the number of consecutive repeat trials before a switch, leading to two conditions of run length: the short runs consisting of 2-3 consecutive repeat trials, and the long runs comprising 5-6 consecutive repeat trials (henceforth, “Run length study”). The second study focused on error monitoring and was designed with the intention of inducing more speech errors through time pressure (henceforth, “Time pressure study”). To achieve this, participants underwent a speed training session before the experiment, with implicit time limits calibrated per participant.

**Figure 1.**
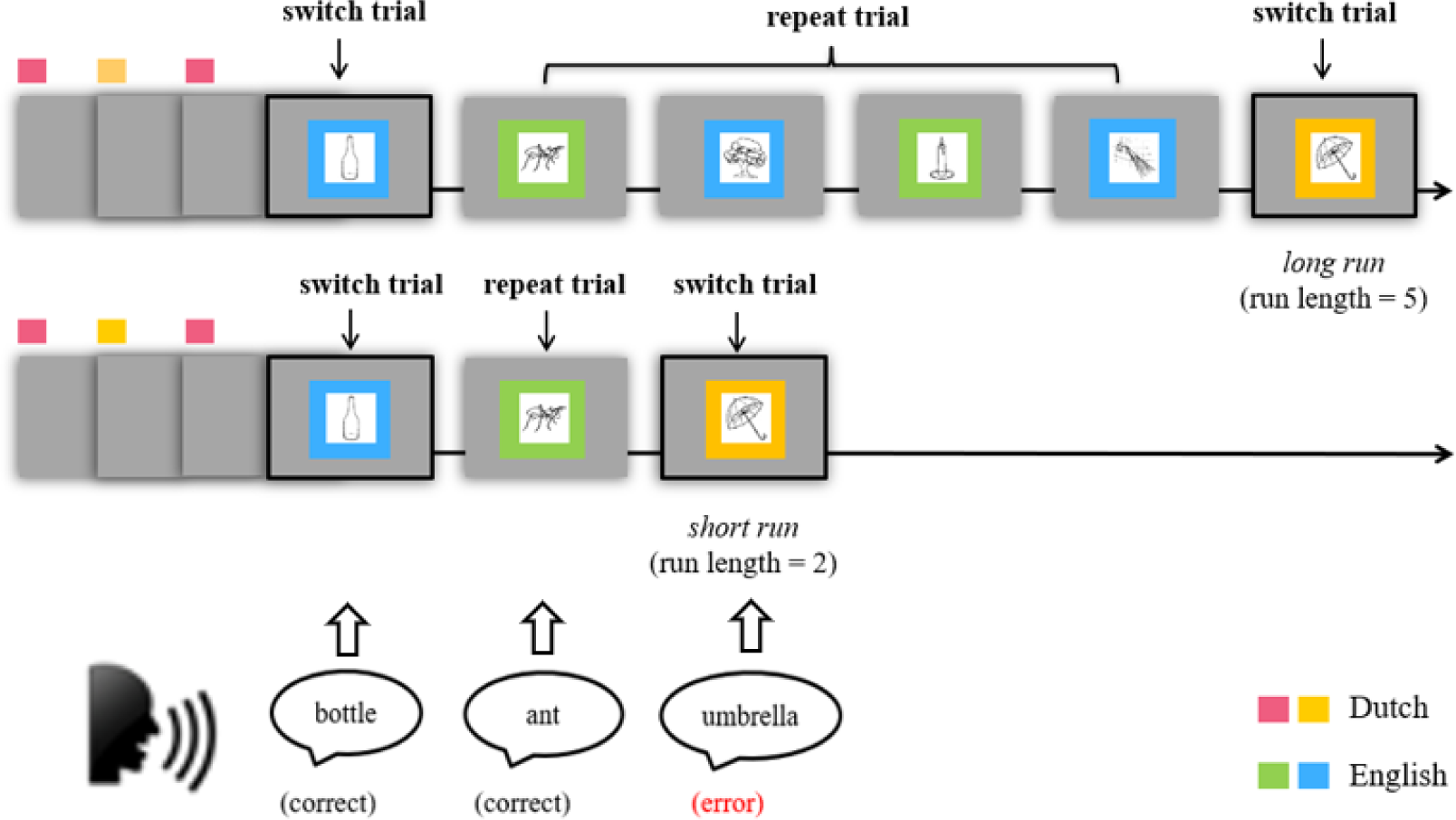
Experimental designs. Participants name pictures in Dutch or in English according to the colour cues. Zheng et al., (2020) manipulated the length of same-language runs before switch trials, creating short runs (2-3 consecutive repeats) and long runs (5-6 consecutive repeats) to investigate inhibitory control. Zheng et al., (2018a) implemented time pressure on participants, deliberately inducing more speech errors, as shown in the rectangular graph.

### EEG Acquisition

A consistent EEG data acquisition protocol was used for both original studies, involving continuous recording of EEG data at 500 Hz with a band-pass filter of 0.016–125 Hz. Fifty-seven active Ag–AgCl ring electrodes were used following the international 10-20 standard. The left mastoid served as the online reference. To facilitate visual inspection, bipolar electrooculograms were recorded with vertical electrodes above and below the right eye, and horizontal electrodes on the left and right temples. Electromyographic activities were recorded with electrodes on the upper lip and throat.

### Data Analysis

Data reanalysis focused on three language control processes: language switching (switch trials vs. repeat trials, performed on data of both studies), inhibitory control (switch trials following short runs vs. switch trials following long runs, performed only on data from the Run length study), and error monitoring (switch trials with language selection errors vs. switch trials with correct responses, Time pressure study only).

#### Behavioural Analysis

In parallel to the contrasts of interest in the EEG analysis, we also reanalysed the behavioural data. In both original studies, participants’ RTs during picture naming were recorded and manually corrected offline. Responses were categorized as correct or speech errors. Speech errors included language-selection errors (e.g., saying the Dutch word “paraplu” instead of the English equivalent “umbrella”) and other types of errors (e.g., self-corrections and disfluencies). RT and error rate were compared for two sets of conditions (switch vs. repeat trials, switches after short runs vs. long runs). RT analyses excluded speech error trials and post-error trials. All the trials at the beginning of each block were also excluded since they do not involve switch or repeat conditions. For inhibitory control conditions (short run vs. long run), all the same-language runs which contain error trials were also excluded because an error response can change the nature of the subsequent trial, for example, converting a long run into a short run. Likewise, error rate analyses excluded the trials at the beginning of each block and post-error trials for both sets of conditions. All the runs with errors were excluded solely for inhibitory control conditions.

Behavioural data underwent statistical analyses using version 1.1.26 of the lmer4 package (Bates et al., 2014) within R software (R Core Team, 2023). We used a generalized linear mixed-effects model of RTs as a function of the corresponding conditions, incorporating random intercepts for both participants and pictures, as well as random slopes of condition for participants and pictures. Only when the model with maximal random effects failed to converge, we simplified the model by removing the interactions and if needed, the main effects in the random structure. An analogous analysis was conducted for error rate using binomial family. The glmer models are provided in the Appendix.

#### EEG Preprocessing

To optimize the analysis of oscillatory data, we re-preprocessed the raw EEG data using the open-source toolbox Fieldtrip (Oostenveld et al., 2011) in MATLAB (R2022a, The Math Works, Inc). Stimulus-locked analysis was employed to explore oscillations linked to language switching (switch vs repeat) and inhibitory control (short run vs long run). Response-locked analysis was utilized to investigate the oscillations associated with error monitoring (language-selection errors vs correct responses). Consequently, stimulus-locked analyses were conducted on both datasets, while response-locked analysis was exclusively performed for the Time pressure study. The data were initially segmented into long, stimulus-locked epochs for analysis, to facilitate the Independent Component Analysis (ICA) process and to include the time points around response onset for the response-locked analysis of the Time pressure study. For Run length study, the epochs spanned from −750 ms to 1500 ms relative to picture onset; for Time pressure study, the epochs encompassed from −750 ms to 2500 ms relative to picture onset, in order to include the response locked data. The data were then re-referenced to linked mastoids and band-pass filtered between 0.1 Hz and 40 Hz. The artifact correction and rejection procedure, done with the experimenter blinded for condition in each study, was as follows. We took an ICA approach to effectively eliminate eye artifacts (e.g., eye blinks and saccades), accompanied by two rounds of additional visual artifact rejection. The first round was performed prior to ICA to discard prominent artifacts that could potentially interfere with the ICA results such as head movements and technical artifacts. Bad channels that exhibited significant disturbances were also excluded from the analysis during this phase. Following the ICA, the second round of artifact rejection was conducted to further eliminate any residual artifacts that might have remained (e.g., muscle artifacts). Prior to the second round of artifact rejection, the cleaned data underwent specific segmentation to cater to stimulus-locked analysis and response-locked analysis (epochs from −750 ms to 1000 ms relative to picture onset or vocal response, respectively). Excluded individual channels were interpolated using neighbouring channels. On average, 2.1% of the stimulus-locked data and 0.4 channels per participant were discarded for data from Run length study; 2.9 % of the stimulus-locked data, 4.6 % of the response-locked data and 1.6 channels per participant were discarded for Time pressure study.

#### EEG Data Analysis

Time-frequency representations (TFRs) were computed for both stimulus-locked data and response-locked data, from −250 ms to 750 ms relative to stimulus and response onset, respectively. The time windows were constrained by the segmentation to ensure integral numbers for the computed frequencies. We used a fixed-length window Hanning taper technique, with a time window of 500 ms, sliding in 2 ms time steps and 2 Hz frequency steps, covering the frequency range of interest from 2 Hz to 12 Hz. All trials excluded from the behavioural analysis were also excluded for EEG data analyses.

#### Statistical Testing

Non-parametric cluster-based permutation tests were conducted to evaluate the difference in theta oscillatory power between conditions (Maris & Oostenveld, 2007). We computed power estimates for each participant under three sets of conditions: switch trials vs. repeat trials, switch trials following short runs vs. those following long runs, and switch trials with language selection errors vs. those with correct responses. Spatial-spectral-temporal data (time, frequency, channel) was generated in three-dimensional spaces. Dependent samples *t*-tests were performed on the spatial-spectral-temporal data points within the theta band (4-8 Hz) to assess mean differences between conditions across 15 frontal central channels (F3, F1, Fz, F2, F4, FC3, FC1, FCz, FC2, FC4, C3, C1, Cz, C2, C4), within the full post-stimuli/response time window of 0-750 ms for both stimulus- and response-locked data. All neighbouring data points that exceeded the cluster-forming threshold (alpha = 0.05) were grouped into clusters, and the *t*-statistics within each cluster were summed into a cluster-based permutation statistic. Next, the Monte Carlo approximation algorithm was used to randomly partition the trial data across conditions 1000 times within participants, and the largest cluster-based *t*-statistics calculated for each random partition were grouped into a Monte Carlo permutation distribution. This permutation distribution was then compared against the calculated cluster-based permutation statistic, with the clusters that exceeded the 2.5th percentile or fell below the 2.5th percentile of the permutation distribution deemed statistically significant.

## Results

### Run Length Study

#### Behavioural Results

**Language switching.** Figure 2 (upper panel) displays the median RTs and error rates for switch vs. repeat trials for Run length study. Participants responded more slowly on switch trials (*Mean* = 1027 ms, *SD* = 279) compared to repeat trials (*Mean* = 930 ms, *SD* = 276), indicating a switch cost effect [*β* = 96.60, *SE* = 2.54, *t* = 37.97, *p* < .001]. Additionally, participants made more errors on switch trials (*Mean* = 6.18%, *SD* = 2.29%) compared to repeat trials (*Mean* = 3.02%, *SD* = 1.36%) [*β* = 0.72, *SE* = 0.13, *z* = 5.57, *p* < .001].

**Figure 2.**
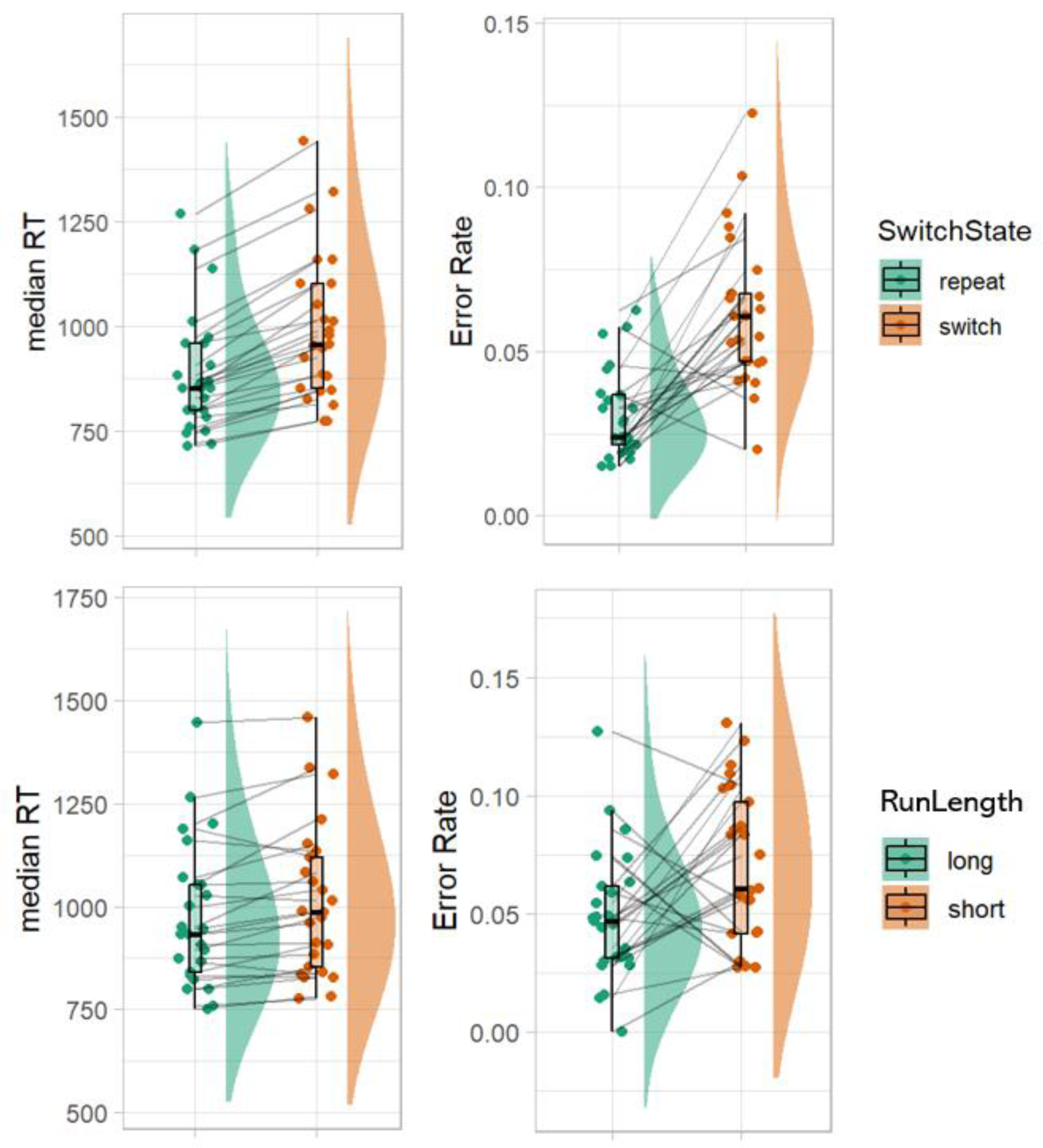
Behavioural results for the Run length study. The upper panel shows the raincloud plots of participants’ error rates and median RTs for language switching (switch vs. repeat). The lower panel presents the raincloud plots of participants’ error rates and median RTs for inhibitory control (switches after a short run vs. long run). The density plot depicts the distribution of data across participants. The top and bottom edges of the box plots correspond to the first and the third quartiles, respectively. The line inside the box represents the median of the group. The individual data points represent the individual medians.

**Inhibitory control.** Figure 2 (lower panel) illustrates the median RTs and error rates for switches after short runs vs. switches after long runs. Participants responded more slowly on switch trials following a short run (*Mean* = 1045, *SD* = 290) as compared to those following a long run (*Mean* = 1005, *SD* = 271) [*β* = 40.74, *SE* = 10.42, *t* = 3.91, *p* < .001]. Descriptively, there were more errors made on switch trials following short runs (*Mean* = 6.96%, *SD* = 3.28%) compared to long runs (*Mean* = 4.87%, *SD* = 2.76%), but the difference was not significant [*β* = 0.30, *SE* = 0.20, *z* = 1.47, *p* = .141].

#### EEG Results

**Language switching.** Figure 3A (left panel) depicts the relative power differences between switch and repeat trials within the 2-12 Hz frequency range, time-locked to the picture onset from −250 to 750 ms, averaged over 15 frontocentral channels. A statistically significant difference between switch and repeat conditions (*p* = .012) was detected between 0***-***750 ms post stimulus in the theta band (4-8 Hz) via a cluster-based permutation test. The effect was most prominent between 100 and 500 ms after the stimulus onset and around 4-6 Hz. Figure 3A (right panel) features a scalp topographical map covering this effect, which is clearly located at the central electrodes.

**Figure 3.**
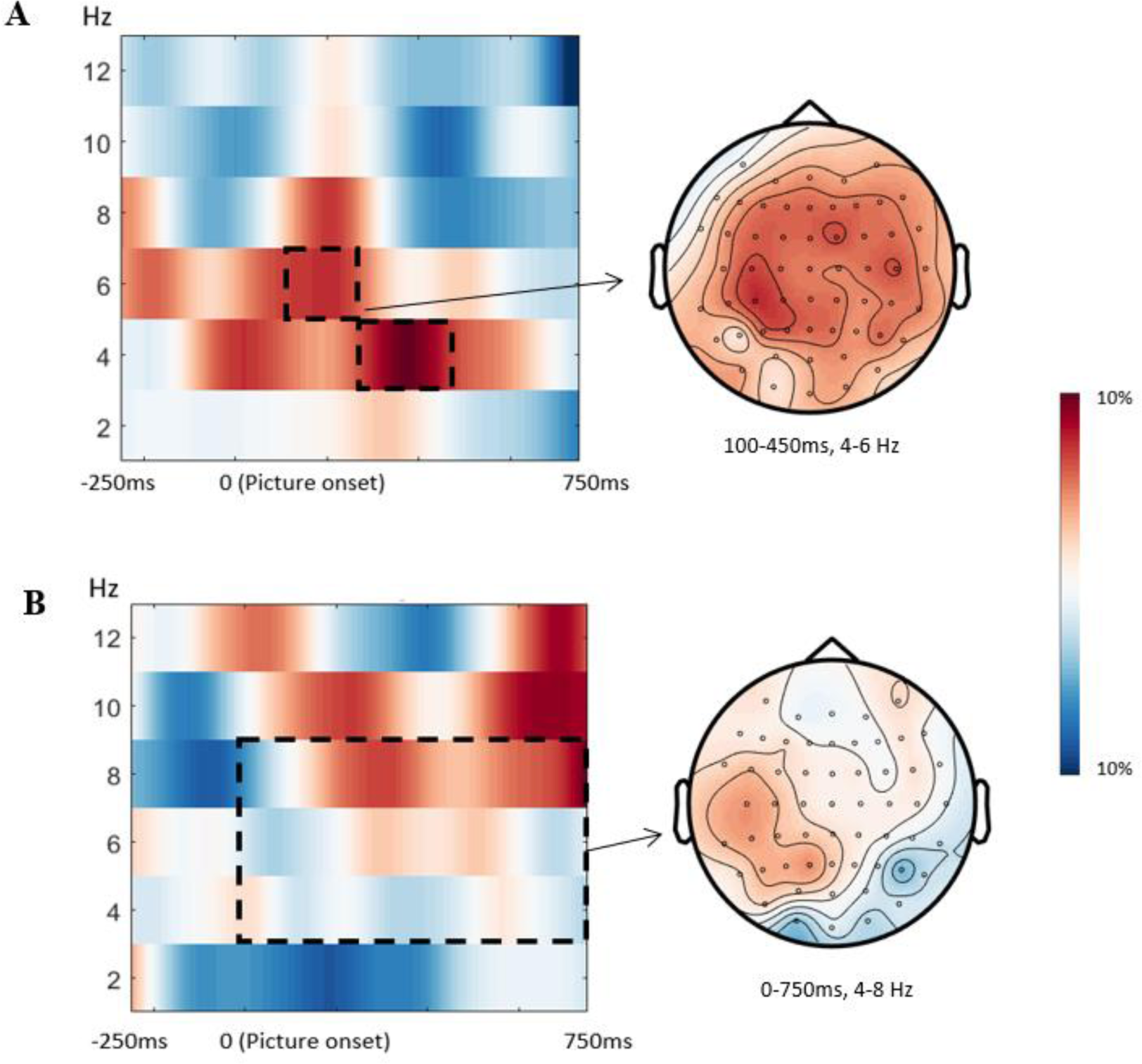
Time-resolved power, in relative difference, for language switching and inhibitory control in the Run length study. **A.** Left: stimulus-locked time-resolved spectrum of the contrast between switch versus repeat trials, averaged over 15 frontocentral channels (F3, F1, Fz, F2, F4, FC3, FC1, FCz, FC2, FC4, C3, C1, Cz, C2, C4). Right: topography of the contrast (switch vs. repeat) in the theta band (4-8 Hz) between 100 and 450 ms post picture onset. **B.** Left: stimulus-locked time-resolved spectrum of the contrast between switch trials following a short run vs. those following a long run over the same cluster of frontocentral channels. Right: topography of the contrast in the theta band (4-8 Hz) between 0 and 750 ms post picture onset, as indicated by the dashed line.

**Inhibitory control.** Figure 3B shows the power difference between switch trials following a short run versus a long run within the 2-12 Hz frequency range, between −250 and 750 ms relative to the picture onset over 15 frontocentral channels. There was no statistically significant difference found between the short and long runs in the theta band (4-8 Hz) within the full post-picture time window (0-750 ms, *p* = .793).

**Figure 4.**
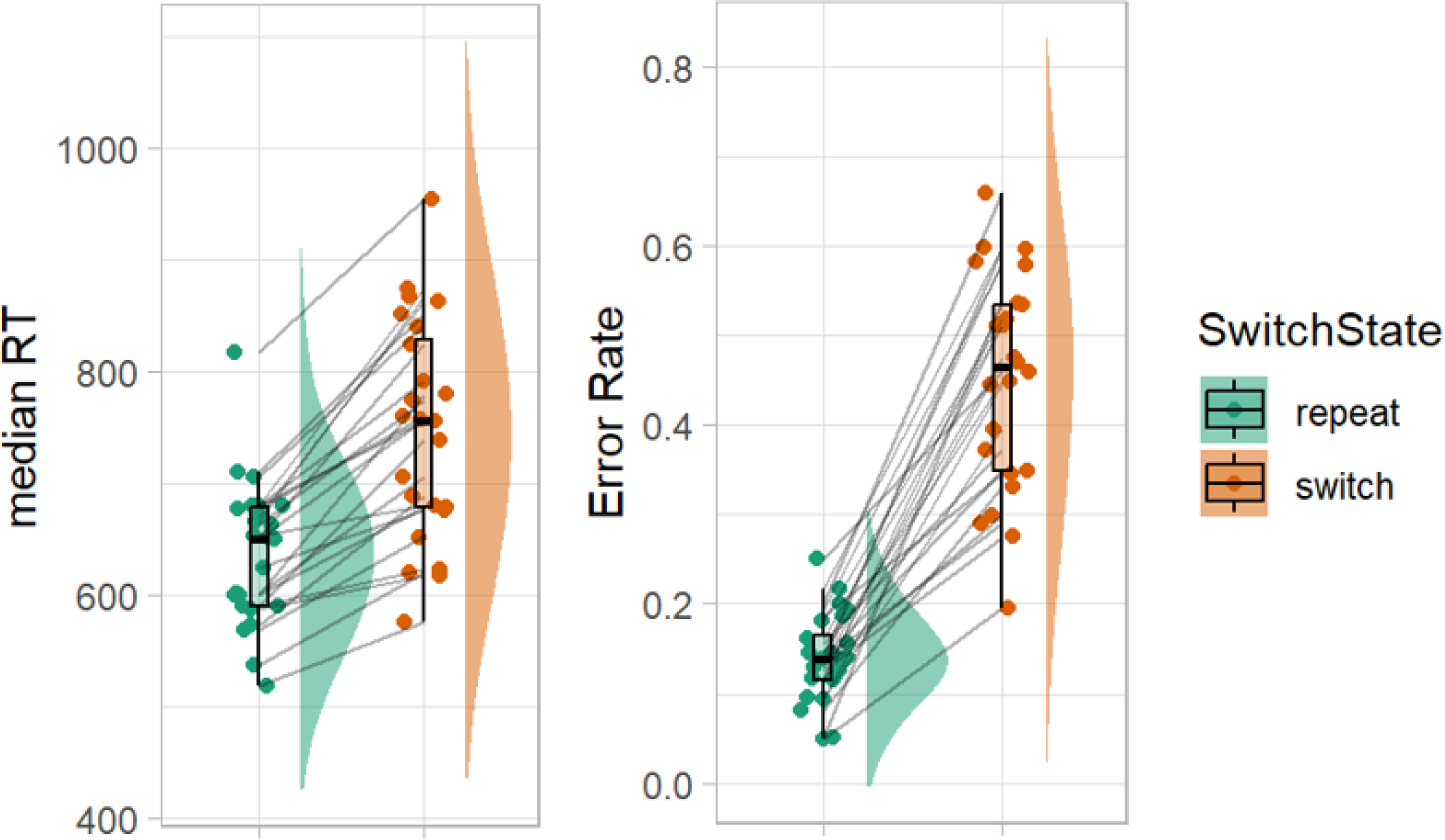
Raincloud plots illustrating the error rates (left) and median RTs (right) of participant picture naming for language switching (switch vs. repeat trials) in the Time pressure study. The density plot depicts the distribution of data across participants. The top and bottom edges of the box plots correspond to the first and the third quartiles, respectively. The line inside the box represents the median of the data. The individual data points represent individual observations.

### Time Pressure Study

#### Behavioural Results

**Language switching.** As shown in Figure 5, similar results were found for picture naming performance, where participants responded more slowly on switch trials (*Mean* = 786 ms, *SD* = 191) compared to repeat trials (*Mean* = 667, *SD* = 142) [*β* = 83.18, *SE* = 3.66, *t* = 22.71, *p* < .001]. The RTs were much shorter in Study 2, since participants named pictures under implicit time pressure. Due to the time pressure manipulation, participants made a substantial number of errors in the naming task (21.92%). More errors were observed on switch trials (*Mean* = 44.45%, *SD* = 12.11%) compared to repeat trials (*Mean* = 13.76%, *SD* = 4.88%) [*β* = 1.62, *SE* = 0.12, *z* = 13.14, *p* < .001]. Out of all the switch trials, 44.45% resulted in errors. Within the error trials, 83.89% were language-selection errors.

**Figure 5.**
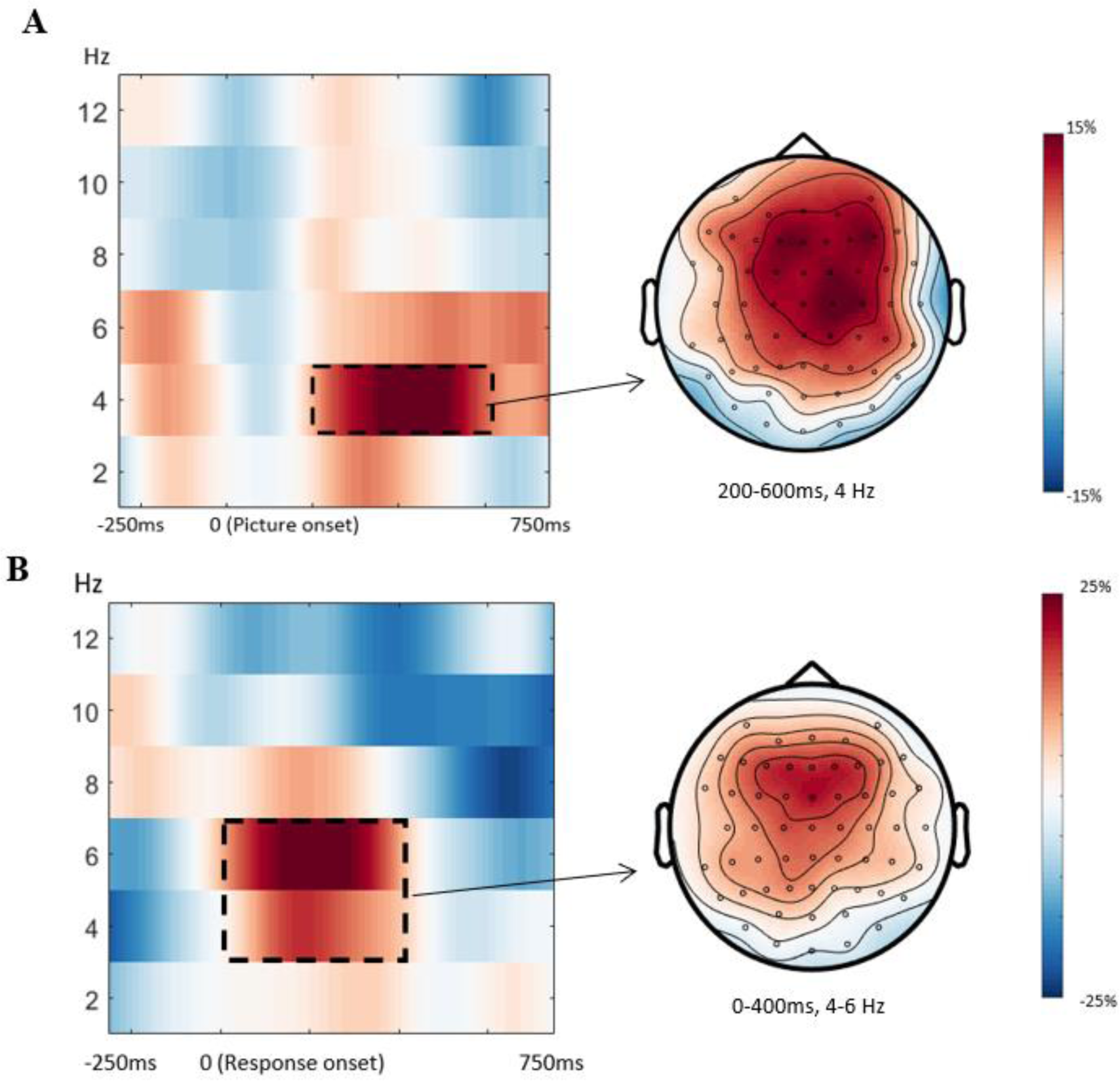
Time-resolved power, in relative difference for language switching and error monitoring conditions in the Time pressure study. **A.** Left panel shows the stimulus-locked time-resolved spectrum of the contrast between switch versus repeat trials, averaged over 15 frontocentral channels (F3, F1, Fz, F2, F4, FC3, FC1, FCz, FC2, FC4, C3, C1, Cz, C2, C4). The right panel shows the topography of the contrast (switch vs. repeat) at 4 Hz between 200 and 600 ms post picture onset (dashed line in the left panel). **B.** The left panel depicts response-locked time-resolved spectrum revealing the contrast between switch trials with language selection errors and switch trials with correct responses over the same cluster of frontocentral channels. The right panel of the figure showcases the topography of the contrast within the frequency range of 4-6 Hz during the time period spanning from 0 to 400 ms after the response onset, as indicated by the dashed line.

**Error monitoring.** Considering error monitoring and RT reflect distinct processes, where error monitoring pertains to error detection whilst RT relates to the efficiency of motor response execution, it would be less informative to directly compare RTs for error monitoring. Instead, we described percentages for error trials relative to total trials and for language-selection errors relative to error trials in both switch and repeat conditions, as shown above.

#### EEG Results

**Language switching.** Similar patterns of power change between switch and repeat trials were observed during language switching as in the Run length study. Figure 6A (left panel) illustrates the power changes within the 2-12 Hz frequency range over frontocentral channels, between −250 ms to 750 ms relative to the picture onset. A statistical analysis confirmed a significant increase in theta power (4-8 Hz) in the frontocentral electrodes between switch and repeat conditions (*p* = .002) from 0 to 750 ms post-stimulus. The effect was most pronounced around 4Hz between 200-600 ms post stimulus. The topographical map in Figure 6A (right panel) reveals that the effect is primarily distributed in the frontocentral region.

**Error monitoring.** The relative power difference between switch trials with language selection errors and switch trials with correct responses is displayed in Figure 6B (left panel), within the 2-12 Hz frequency range and time-locked to the response onset from −250 ms to 750 ms, averaged over frontocentral channels. A statistically significant theta power increase was detected in switch trials with language selection errors compared to those with correct responses (*p* = .006), within the time window of post response onset. The effect exhibited its greatest prominence within the 4-6 Hz frequency range, commencing from the moment of the response and extending up to 400 ms thereafter. The scalp topographical map presented in Figure 6B (right panel) demonstrates that the effect primarily manifests in the frontocentral channels.

## Discussion

To explore the role of domain-general cognitive control mechanisms in language switching, this study investigated midfrontal theta oscillations, an index of cognitive control, in bilingual word production. Specifically, we focused on three fundamental control processes: language switching, inhibitory control, and error monitoring, by reanalysing the EEG data from two previous studies (Zheng et al., 2020; Zheng et al., 2018a).

In both studies, participants named pictures in English or in Dutch according to a colour cue. We observed the established switch-cost effect, where participants made slower responses and exhibited more errors on switch trials compared to repeat trials (Christoffels et al., 2007a; Costa & Santesteban, 2004; Jackson et al., 2001; Verhoef et al., 2009; Zheng et al., 2020). This suggests increased top-down control at switch, potentially arising from the necessity for task reconfiguration to adjust to the new language context or the persistent inhibition that needs to be overcome when switching (Green, 1998; Monsell, 2003). In line with the behavioural results, we also observed consistent modulation of midfrontal theta power as a function of language switching: switch trials exhibited midfrontal theta power increase compared to repeat trials, with this enhancement being predominantly distributed in the frontal-central scalp region. Given the similar pattern of midfrontal theta effects reported in non-linguistic task switching (Cooper et al., 2019), it suggests that language and task switching operate using common mechanisms. This finding supports prior behavioural research showing the similar switch-cost effect in linguistic and non-linguistic tasks (Declerck et al., 2017; Prior & Gollan, 2013; Timmermeister et al., 2020). It also aligns with fMRI studies revealing shared neural circuits during language and task switching (e.g., Abutalebi & Green, 2008; de Bruin et al., 2014; Guo et al., 2011; Wang et al., 2009), along with ERP studies reporting heightened N2 effects during language and task switches compared to repeats (Jackson et al., 2001; Kang et al., 2020; Verhoef et al., 2010; Zheng et al., 2020). Consequently, our results contribute to the growing empirical evidence supporting the involvement of domain-general cognitive control mechanisms in the language switching process.

Likewise, we observed an increase in midfrontal theta power during switch trials where participants made language selection errors compared to those with correct responses, suggesting that more top-down control was involved following error commission. This enhanced control can arise from the error monitoring process which demands greater mental effort to resolve the conflict between the erroneously executed response and the intended correct response (Yeung et al., 2004). Furthermore, it can be linked to the idea that the monitoring system leverages errors as valuable inputs for feedback processing and reinforcement learning, which necessitates a higher level of control (Brown & Braver, 2005; Zheng et al., 2018a). The similar midfrontal theta modulation in response to error commission in both action monitoring (as observed by Cavanagh et al., 2012 and Luu et al., 2004) and speech production monitoring suggests the presence of common underlying mechanisms. Our finding further substantiates previous neuroimaging studies highlighting the shared post-error ERN component (Acheson et al., 2012; Coulter & Phillips, 2022; Zheng et al., 2018a) and increased activity in the ACC for action and speech monitoring (Abutalebi et al., 2012; Christoffels et al., 2007b; Gauvin et al., 2016). Taken together, our results support the notion that speech monitoring is not an exceptional case but rather a part of domain-general action monitoring processes.

Besides our investigation of language switching and speech monitoring, we also found the established run-length effect behaviourally, where bilinguals responded more slowly in switch trials following a short sequence of repeat trials as compared to those following a long sequence of repeat trials. This reflects the fact that inhibition of the non-target language diminishes over the repeated use of the same language, which leads to a less demanding language selection at switch and thus decreased involvement of top-down control (Zheng et al., 2020). Contrary to the behavioural data, however, we failed to observe midfronal theta power modulation as a function of run length: there was no theta power difference between short-run switches and long-run switches. The absence of this midfrontal theta effect related to overcoming inhibition during bilingual switching highlights the divergent neurophysiological pattern observed in speech production as compared to non-linguistic inhibitory control (Eisma et al., 2021; Nigbur et al., 2011). This finding is contrary to prior fMRI studies revealing the shared neural circuits associated with inhibition across domains (de Bruin et al., 2014; Rossi et al., 2021), thus failing to support the proposal of activation of domain-general inhibitory control process in bilingual switching.

One potential explanation for this missing theta effect is that inhibition in language control operates through a distinct mechanism that differs from the traditional cognitive control processes typically associated with the prefrontal network. Similarly, Branzi et al.’s (2016) also reported a dissociation in inhibitory control between language switching and non-language switching tasks. This might also explain why some studies have not consistently observed bilingual advantages in inhibitory control tasks (Paap & Greenberg, 2013; Von Bastian et al., 2016). Besides the assumption that inhibitory control operates differently in action control and language control, the lack of midfrontal theta oscillations in relation to the behavioural run-length effect could also imply that the run-length effect is not primarily driven by the inhibition of the non-target language but the enhancement of the target language (Allport & Wylie, 1999; Philipp et al., 2007). As individuals repeatedly use the same language, the enhancement of the target language diminishes, resulting in a reduced need for control when switching to another language. However, as also asserted by Zheng et al. (2020), we believe that whether inhibition or enhancement is at play, both mechanisms still reflect a form of top-down control. Thus, if midfrontal theta reflects general-purpose cognitive control processes, it should also reflect enhancement. Furthermore, the divergent findings pertaining to the N2 component and midfrontal theta oscillations observed in the same dataset of Zheng et al. (2020), both of which serve as neurophysiological markers for inhibitory control, lead us to postulate that these markers reflect distinct neural processes (Cohen & Donner, 2013). Indeed, studies in non-linguistic domains have reported limited and nonsignificant associations between ERP and oscillatory measures linked to conflict resolution (Cavanagh et al., 2012) and feedback processing (Cohen et al., 2007). Thus, ERP and neural oscillation analyses appear to offer complementary insights into language control processes. Furthermore, the midfrontal theta effect we observed on the scalp level may not represent a unified cognitive control mechanism but could potentially originate from various neural networks (Messel et al., 2021; Zuure et al., 2020). Utilizing both fMRI and EEG techniques in future studies would be advantageous for understanding shared or distinct neural mechanisms distinguished by midfrontal theta activity during language control and action control.

In summary, our study largely supports the involvement of domain-general cognitive control processes in bilingual production, through midfrontal theta oscillations in both language switching and speech monitoring processes (but not in inhibitory control). This neural marker observed in our data closely resembles those reported in executive functions across all dimensions (i.e., spatial, temporal, and frequency). As one of the first attempts to explore midfrontal theta oscillations in bilingual switching, the current study offers a novel perspective on the question of whether language control is entwined with the broader domain-general cognitive control processes. Understanding the connection between two types of control encourages collaboration between the two distinct research traditions, thus contributing to the development of a unified theoretical framework for understanding the human brain and cognition. In our pursuit of understanding the nature of language control, our study expands the empirical investigation of brain oscillations beyond monolingualism to bilingualism. By simultaneously examining the three key processes for both non-linguistic cognitive and language control, our study also provides a more coherent investigation than previous studies targeting these processes in isolation.

## Acknowledgement

We thank Robert Oostenveld and Britta Westner for their advice on EEG data analysis, and Sander Nieuwenhuis for his comments on an early version of the manuscript.

## Data and Code Availability

Data and codes of the original study are available on the Donders Repository (Run Length Study https://data.ru.nl/collections/di/dcc/DSC_2017.00138_605?1; Time Pressure Study: https://data.ru.nl/collections/di/dcc/DSC_2017.00049_995?0). Additional codes for the current follow-up study are available at the Open Science Framework (https://osf.io/yqsw6/).

## Ethics Approval Statement

Both original studies were conducted in accordance with the Declaration of Helsinki, and were approved by the local ethics committee (Faculty Ethics Committee, Radboud University, ECSW 2015-2311-349). The current study reanalysed the anonymized behavioural and EEG data which were publicly shared on the Donders repository.

### Appendix

#### Linear Mixed Effect Models

##### Run Length Study

###### language switching

~~~
# RT for switch vs repeat
glmer.runlength.RT.all <-glmer(RT_corrected ∼ Runlength + (1 | pNumber) + (1 + Runlength | PicNam), data = mydata.runlength.RT.all, family = Gamma(link = “identity”), control = glmerControl(optimizer = ‘bobyqa’))
~~~

~~~
# error rate for switch vs repeat
glmer.swicost.error.all <-glmer(Error ∼ SwitchState + (1 + SwitchState | pNumber) + (1 + SwitchState | PicNam), data = mydata.swicost.error.all, family = binomial)
~~~

###### inhibitory control

~~~
# RT for switches after a short run vs a long run
glmer.runlength.RT.all <-glmer(RT_corrected ∼ Runlength + (1 + Runlength | pNumber) + (1 + Runlength | PicNam), data = mydata.runlength.RT.all, family = Gamma(link = “identity”), control = glmerControl(optimizer = ‘bobyqa’))
# full model fails to converge
~~~

~~~
# error rate for switches after a short run vs a long run
glmer.runlength.error.all <-glmer(Error ∼ Runlength + (1 + Runlength | pNumber) + (1 + Runlength| PicNam), data = mydata.runlength.error.all, family = binomial)
~~~

#### Time Pressure Study

##### language switching

~~~
# RT for switch vs repeat
glmer.swicost.RT.all <-glmer(RT_corrected ∼ SwitchState + (1 + SwitchState | pNumber) + (1 + SwitchState| PicNam), data = mydata.swicost.RT.all, family = Gamma(link = “identity”), control = glmerControl(optimizer = ‘bobyqa’))
~~~

~~~
# error rate for switch vs repeat
glmer.swicost.error.all <-glmer(Error ∼ SwitchState + (1 + SwitchState | pNumber) + (1 + SwitchState| PicNam), data = mydata.swicost.error.all, family = binomial)
~~~

